# Bat coronavirus phylogeography in the Western Indian Ocean

**DOI:** 10.1101/742866

**Authors:** Léa Joffrin, Steven M. Goodman, David A. Wilkinson, Beza Ramasindrazana, Erwan Lagadec, Yann Gomard, Gildas Le Minter, Andréa Dos Santos, M. Corrie Schoeman, Rajendraprasad Sookhareea, Pablo Tortosa, Simon Julienne, Eduardo S. Gudo, Patrick Mavingui, Camille Lebarbenchon

## Abstract

Bats provide key ecosystem services such as crop pest regulation, pollination, seed dispersal, and soil fertilization. Bats are also major hosts for biological agents responsible for zoonoses, such as coronaviruses (CoVs). The islands of the Western Indian Ocean are identified as a major biodiversity hotspot, with more than 50 bat species. In this study, we tested 1,013 bats belonging to 36 species from Mozambique, Madagascar, Mauritius, Mayotte, Reunion Island and Seychelles, based on molecular screening and partial sequencing of the RNA-dependent RNA polymerase gene. In total, 88 bats (8.7%) tested positive for coronaviruses, with higher prevalence in Mozambican bats (20.5% ± 4.9%) as compared to those sampled on islands (4.5% ± 1.5%). Phylogenetic analyses revealed a large diversity of α- and β-CoVs and a strong signal of co-evolution between CoVs and their bat host species, with limited evidence for host-switching, except for bat species sharing day roost sites.

**Importance:** This is the first study to report the presence of coronaviruses (CoVs) in bats in Mayotte, Mozambique and Reunion Island, and in insectivorous bats in Madagascar. Eight percent of the tested bats were positive for CoVs, with higher prevalence in continental Africa than on islands. A high genetic diversity of α- and β-CoVs was found, with strong association between bat host and virus phylogenies, supporting a long history of co-evolution between bats and their associated CoVs in the Western Indian Ocean. These results highlight that strong variation between islands does exist and is associated with the composition of the bat species community on each island. Future studies should investigate whether CoVs detected in these bats have a potential for spillover in other hosts.

## Introduction

The burden of emerging infectious diseases has significantly increased over the last decades and is recognized as a major global health concern. In 2018, the World Health Organization (WHO) established the “Blueprint priority disease list”, identifying viruses such as Ebola, Lassa fever, Middle East Respiratory Syndrome (MERS), and Nipah fever as significant threats to international biosecurity ^1^. This list also highlights the potential pandemic risk from the emergence of currently unknown zoonotic pathogens, collectively referring to these unknown threats as “disease X” ^1^. Investigation of the potential zoonotic pathogens in wild animals, particularly vertebrates, is thus critical for emerging infectious disease preparedness and responses.

Bats represent nearly 1,400 species and live on all continents except Antarctica ^2^. They provide key ecosystem services such as crop pest regulation, pollination, seed dispersal, and soil fertilization ^3–10^. Bats are also recognized as reservoirs of many zoonotic pathogens, including coronaviruses (CoVs) ^11–13^. Indeed, several CoVs originating from bats have emerged in humans and livestock with sometimes major impacts to public health. For instance, in 2003, the Severe Acute Respiratory Syndrome (SARS) CoV emerged in humans, after spillover from bats to civets^14–18^, and led to the infection of 8,096 people and 774 deaths in less than a year ^19^.

Our study area spans geographic locations across the islands of the Western Indian Ocean and southeastern continental Africa (SECA) (Figure 1). These islands have diverse geological origins that have influenced the process of bat colonization and species distributions ^20^. The ecological settings and species diversity on these islands for bats are notably different. On Madagascar, more than 45 bat species are known to occur, of which more than 80 % are endemic to the island ^21–23^. The smaller studied islands of the Western Indian Ocean, Mauritius, Mayotte, Reunion Island, and Mahé (Seychelles), host reduced bat species diversity (e.g. three species on Reunion Island), whereas SECA supports a wide range of bat species. To date, several studies have identified bat-infecting CoVs in countries of continental Africa, including Zimbabwe ^24^, South Africa ^25,26^, Namibia ^27^, and Kenya ^28,29^. CoVs have also been reported in fruit bats (Pteropodidae) in Madagascar, where β-coronaviruses belonging to the D-subgroup were identified in *Eidolon dupreanum* and *Pteropus rufus* ^30^.

**Figure 1.**
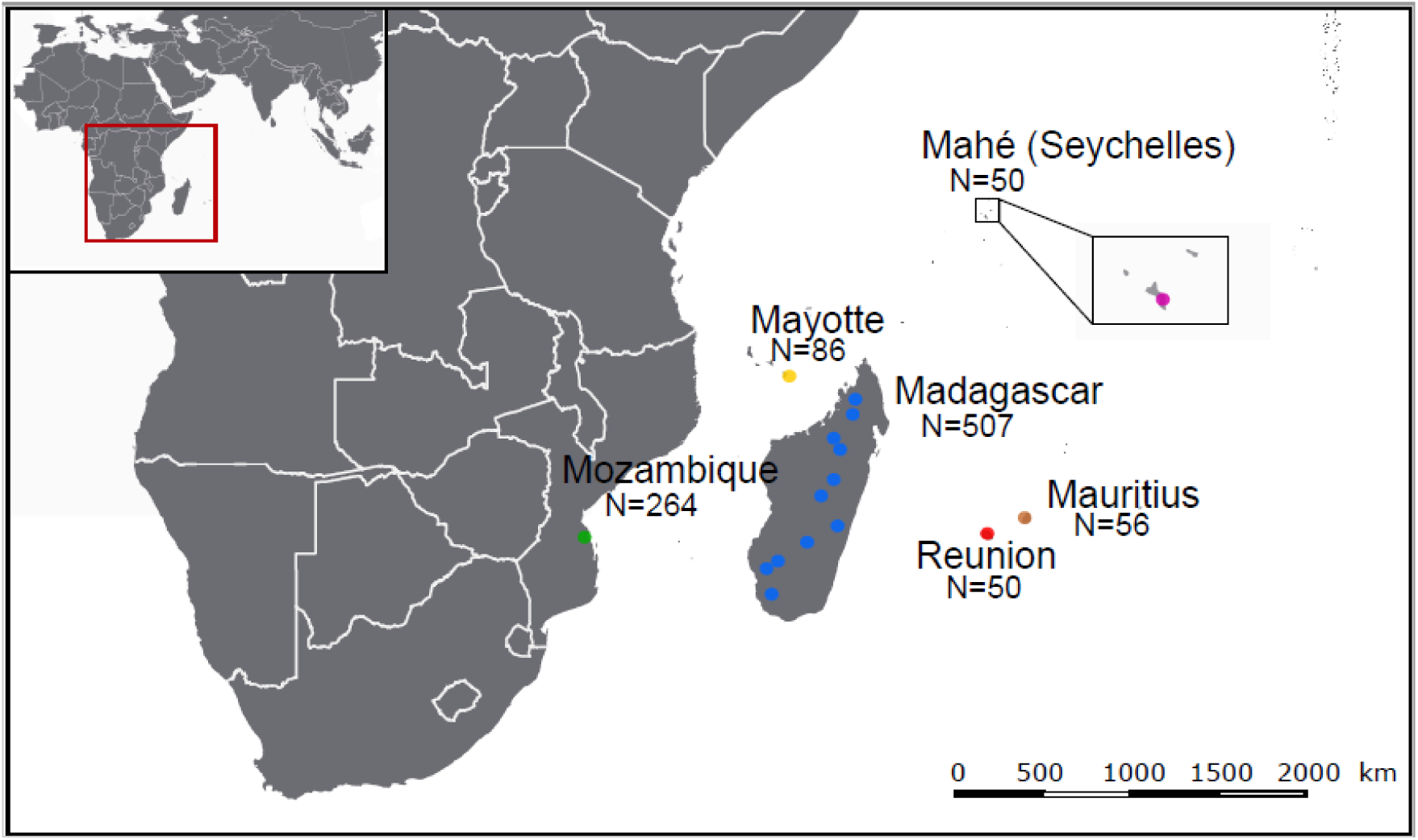
Geographic distribution of the tested samples. N: number of bats sampled for each location. The open-source GIS software, QGIS v.3.6.1, was used to generate the map. http://qgis.osgeo.org (2019).

In this study, we investigated the presence of CoVs in over 1,000 individual bats belonging to 36 species, sampled on five islands (Madagascar, Mauritius, Mayotte, Reunion Island, and Mahé) and one continental area (Mozambique). Based on molecular screening and partial sequencing of the RNA-dependent RNA polymerase gene, we (i) estimated CoV prevalence in the regional bat populations, (ii) assessed CoV genetic diversity, and (iii) identified associations between bat families and CoVs, as well as potential evolutionary drivers leading to these associations.

## Results

### Prevalence of CoV

A total of 1,013 bats were tested from Mozambique, Mayotte, Reunion Island, Seychelles, Mauritius and Madagascar (Figure 1). In total, 88 of the 1,013 bat samples tested positive for CoV by Real-Time PCR (mean detection rate: 8.7%). The prevalence of positive bats was different according to the sampling locations (χ^2^ = 77.0, df = 5; p<0.001), with a higher prevalence in Mozambique (± 95% confidence interval: 20.5% ± 4.9%) than on all Western Indian Ocean islands (4.5% ± 1.5%) (Figure 2). A significant difference in the prevalence of positive bats was also detected between families (χ^2^ = 44.8, df = 8; p<0.001; Supplementary Figure S1). The highest prevalence were observed in the families Nycteridae (28.6 % ± 23.6%) and Rhinolophidae (26.2% ± 11.0%). Bat species had a significant effect on the probability of CoVs detection (χ^2^ = 147.9, df = 39; p<0.001; Supplementary Figure S2). Finally, the prevalence of CoV positive bats in Mozambique was significantly different (N = 264, χ^2^= 22.8, df = 1; p<0.001; Supplementary Figure S3) between February (37.4% ± 9.9%) and May (11.6% ± 4.8).

**Figure 2.**
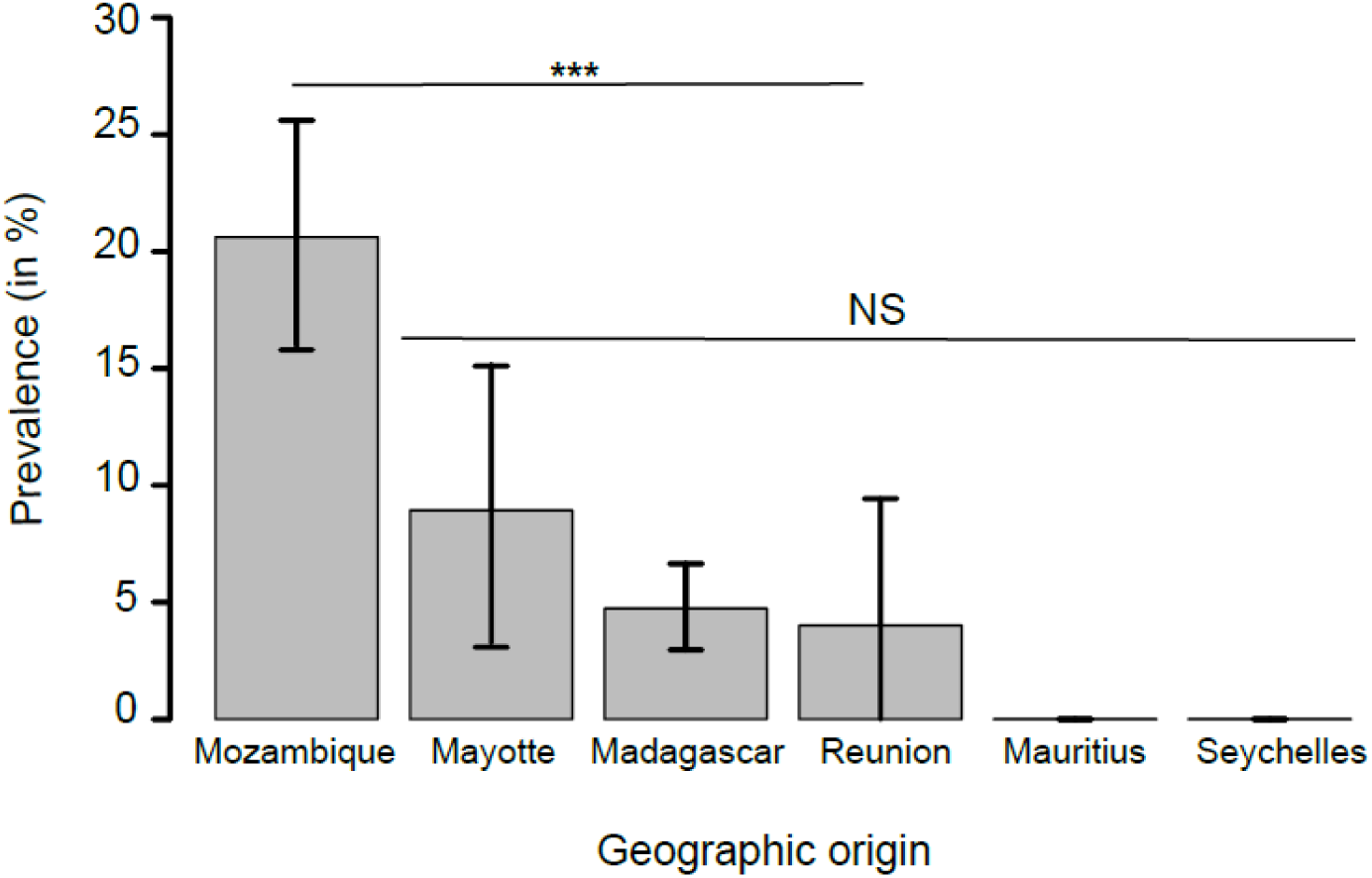
Mean CoV prevalence (± 95% confidence interval) in bats in the Western Indian Ocean. Pairwise test; ***: p<0.001; NS: p>0.05, not significant.

### RdRp sequence diversity

Of the 88 positive samples, we obtained 77 partial RdRp sequences using the Real-Time PCR detection system (179 bp) and 51 longer partial RdRp sequences using a second PCR system (440 bp). Sequences generated with the second system were subsequently used for phylogenetic analyses. Details of the sequenced CoV-positive samples are provided in Supplementary Table S1. Pairwise comparison of these 51 sequences revealed 28 unique sequences, and sequences similarities ranging from 60.2% to 99.8%. The lowest sequence similarity was found in Mozambique (60.2% to 99.8%), then in Madagascar (64.0% to 99.8%). No genetic variation was observed for samples from Mayotte and Reunion Island.

### Phylogenetic structure of CoVs

Sequence comparison indicated that Western Indian Ocean bats harbor a high diversity of both α and β-CoVs, with conserved groups clustering mostly by bat family (Figure 3). Specifically, 25 sequences were identified as α-CoVs, and three sequences were genetically related to the β-CoVs. For α-CoVs, all sequences detected in our tested Molossidae formed a highly supported monophyletic group, including CoV sequences from Molossidae bats previously detected in continental Africa (Figure 4). CoVs detected in *Mops condylurus* (Mozambique), *Mormopterus francoismoutoui* (Reunion Island), *Chaerephon pusillus* and *Chaerephon* sp. (Mayotte), and *Mormopterus jugularis* (Madagascar) shared 90% - 98% nucleotide similarity with a CoV detected in *Chaerephon* sp. in Kenya (Supplementary Table S2). All CoVs found in Miniopteridae clustered in a monophyletic group, including Miniopteridae CoVs sequences from Africa, Asia, and Oceania (Supplementary Table S2). The great majority of α-CoVs detected in Rhinolophidae bats clustered in two monophyletic groups (Figure 3); one with African Rhinolophidae CoVs and one with Asian Rhinolophidae CoVs. We also detected one CoV from *Rhinolophus rhodesiae*, which was 100% similar to a Miniopteridae CoV from this study. Rhinonycteridae CoVs formed a single monophyletic group with NL63 Human CoVs. The Rhinonycteridae CoVs detected clustered with NL63-related bat sequences found in *Triaenops afer* in Kenya (Figure 5) and showed 85% similarity to NL63 Human CoVs (Supplementary Table S2). Hipposideridae α-CoVs mainly clustered into a single monophyletic group, including 229E Human CoV-related bat sequence found in *Hipposideros vittatus* from Kenya (Figure 6; Supplementary Table S2).

**Figure 3.**
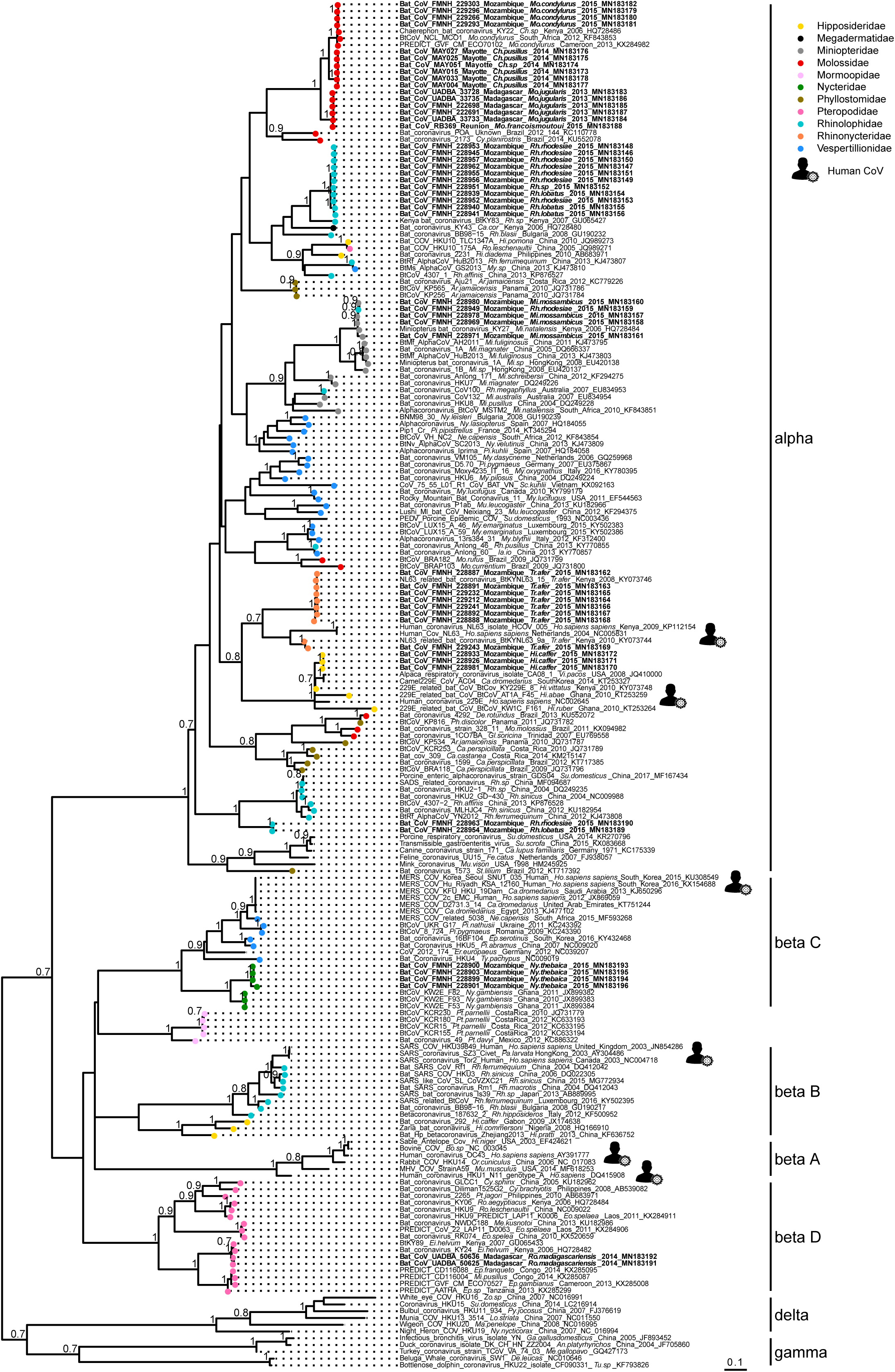
Maximum Likelihood (ML) consensus tree derived from 202 coronavirus (CoV) RNA-dependent RNA-polymerase partial nucleotide sequences (393 bp). Colored circles at the end of branches indicate bat family origin. Sequences in bold refer to bat CoVs detected in this study. Bootstrap values >0.7 are indicated on the tree. Scale bar indicates mean number of nucleotide substitutions per site. The tree was generated with the General Time Reversible evolutionary model (GTR+I+Г, I = 0.18, α = 0.64) and 1,000 bootstrap replicates.

**Figure 4.**
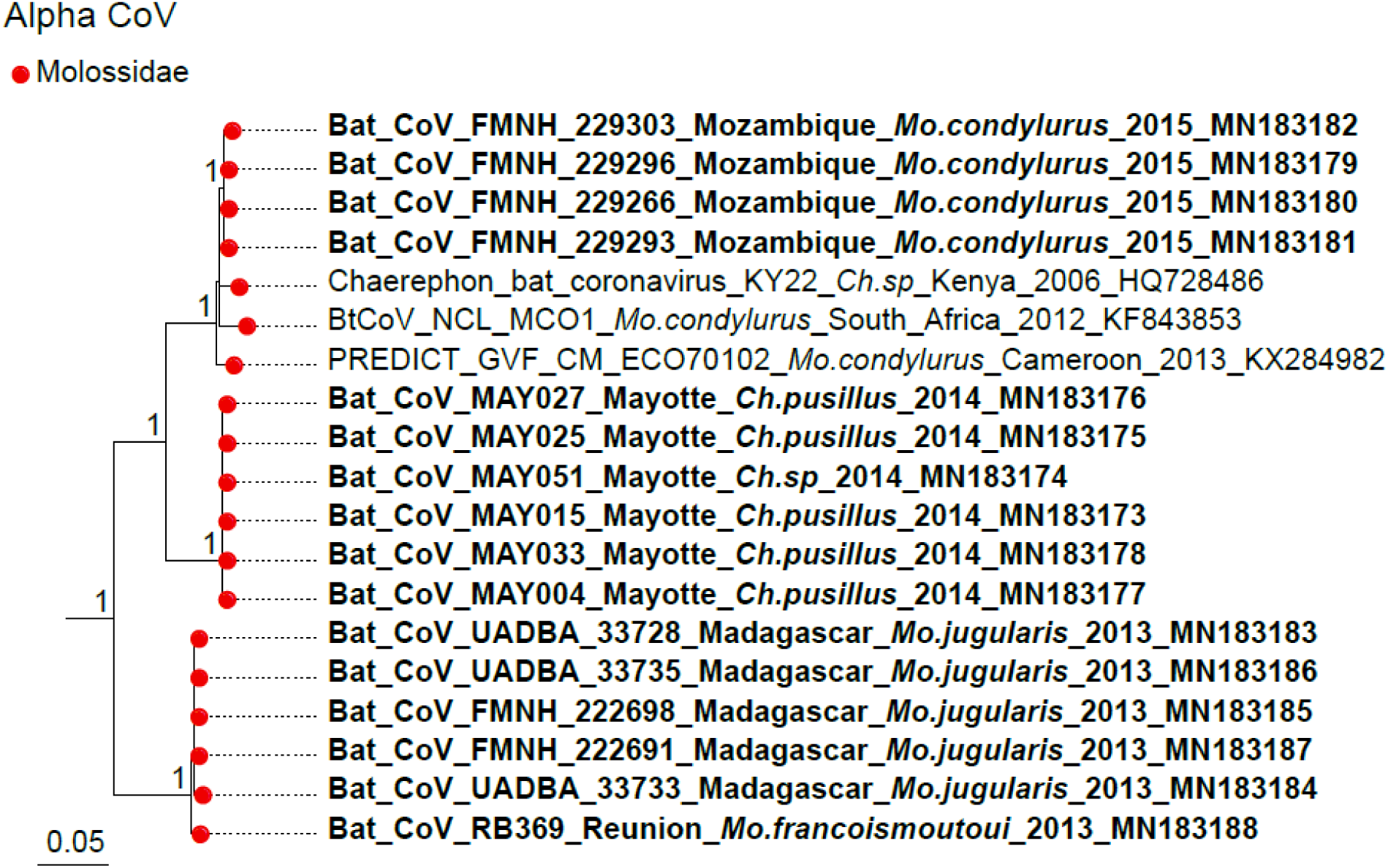
Detail of the α-CoV clade. Molossidae CoVs generated in the study are indicated in bold. This sub-tree is a zoom on Molossidae CoV clade from the tree depicted in Figure 3. Bootstrap values >0.7 are indicated on the tree. Scale bar indicates mean number of nucleotide substitutions per site.

**Figure 5.**
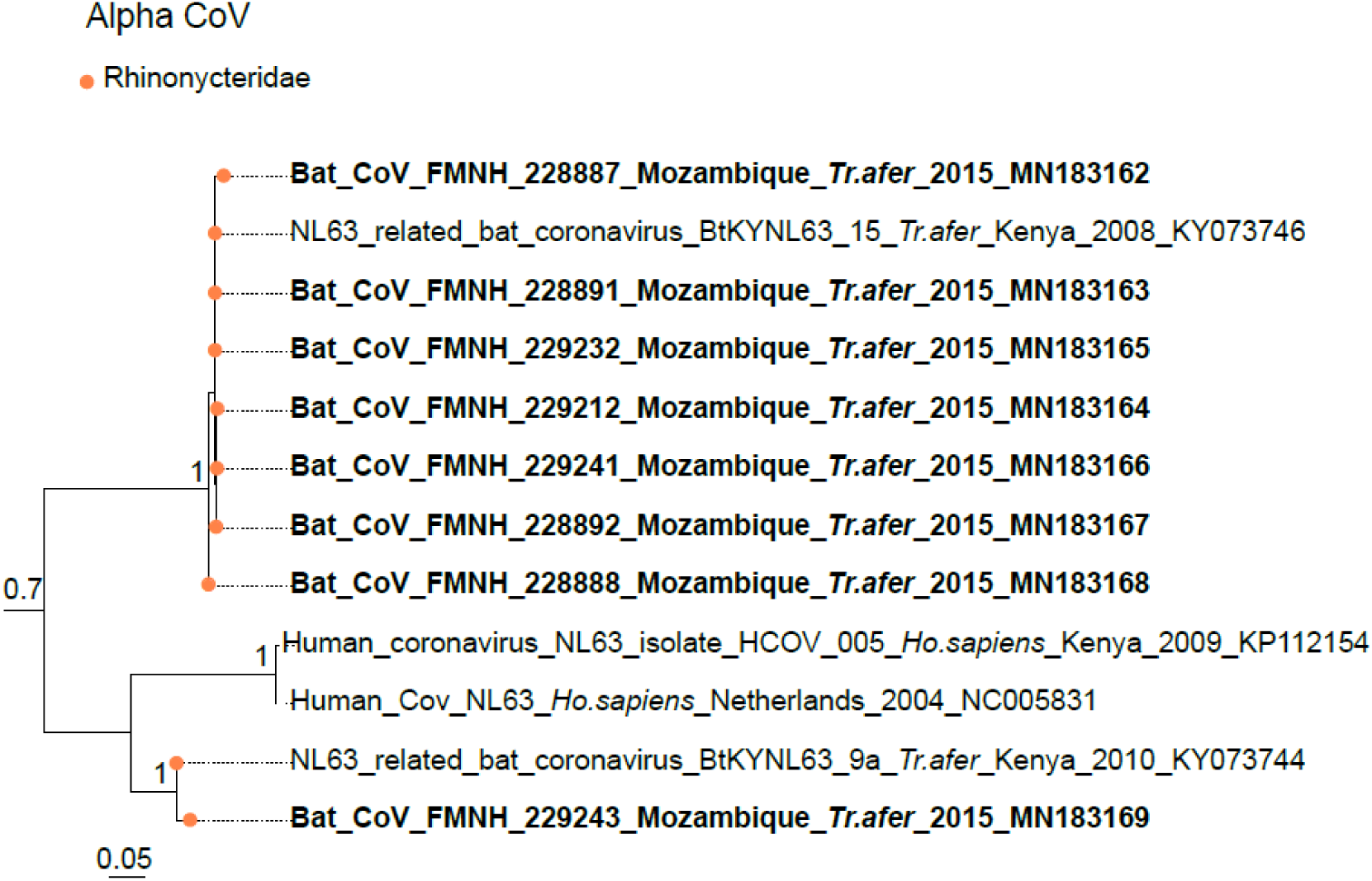
Detail of the α-CoV clade. NL63-like CoVs generated in the study are indicated in bold. This sub-tree is a zoom on NL63 CoV clade from the tree depicted in Figure 3. Only bootstrap values >0.7 are indicated on the tree. Scale bar indicates mean number of nucleotide substitutions per site.

**Figure 6.**
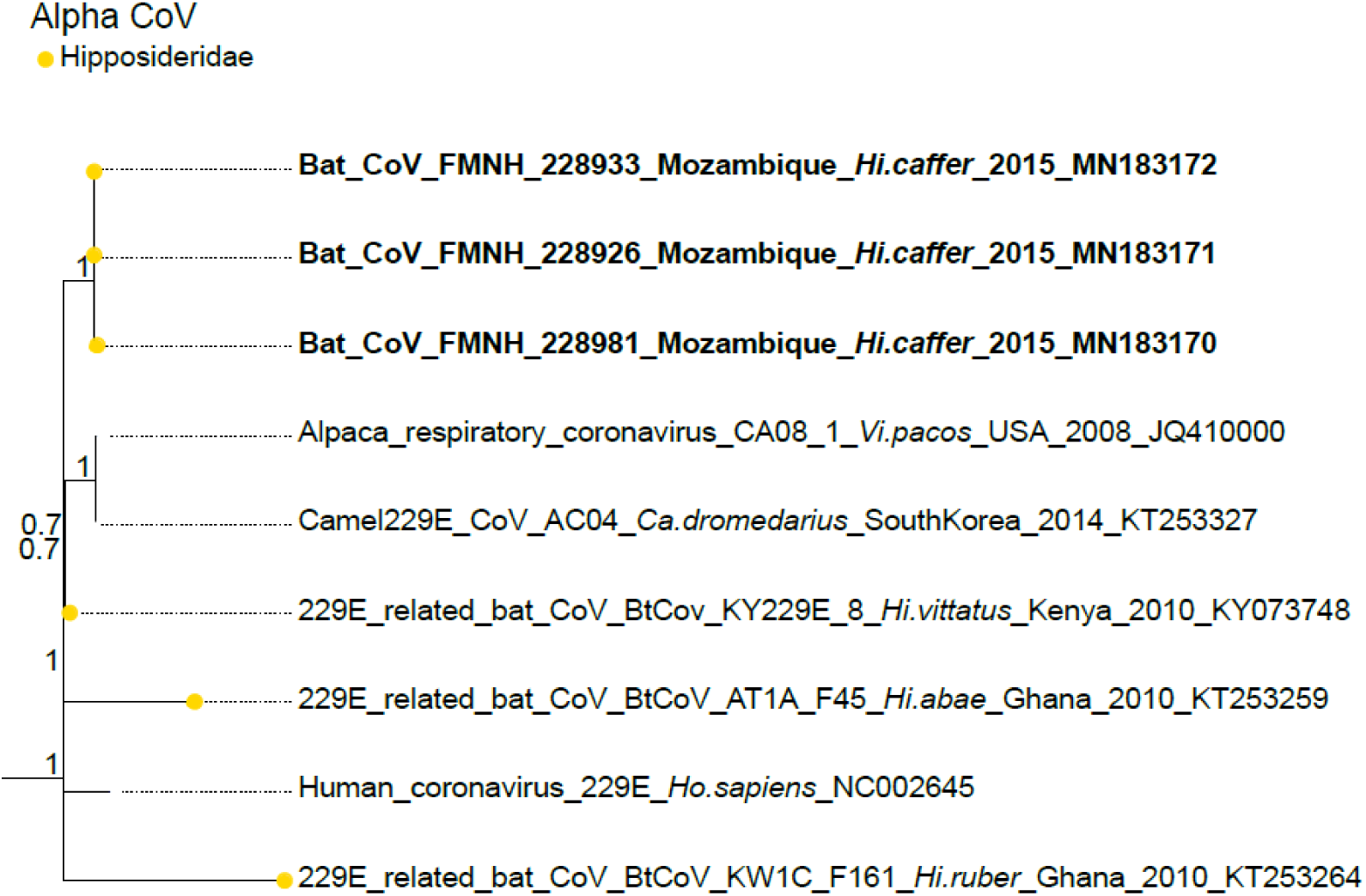
Detail of the α-CoV clade. 229E-like CoVs generated in the study are indicated in bold. This sub-tree is a zoom on NL63 CoV clade from the tree depicted in Figure 3. Bootstrap values >0.7 are indicated on the tree. Scale bar indicates mean number of nucleotide substitutions per site.

For β-CoVs, two sequences obtained from *Nycteris thebaica* clustered in the C-subgroup together with other CoVs previously reported in African *Nycteris sp.* bats (Figure 7). The sequences showed 88% nucleotide identity to a β-C CoV found in *Nycteris gambiensis* in Ghana (Supplementary Table S2). *Rousettus madagascariensis* CoVs clustered with Pteropodidae CoVs belonging to the D-subgroup of β-CoVs (Figure 8). BLAST queries against the NCBI database showed 98% nucleotide identity between CoV sequences from *Rousettus madagascariensis* and a β-D CoV sequence detected in *Eidolon helvum* from Kenya (Supplementary Table S2).

**Figure 7.**
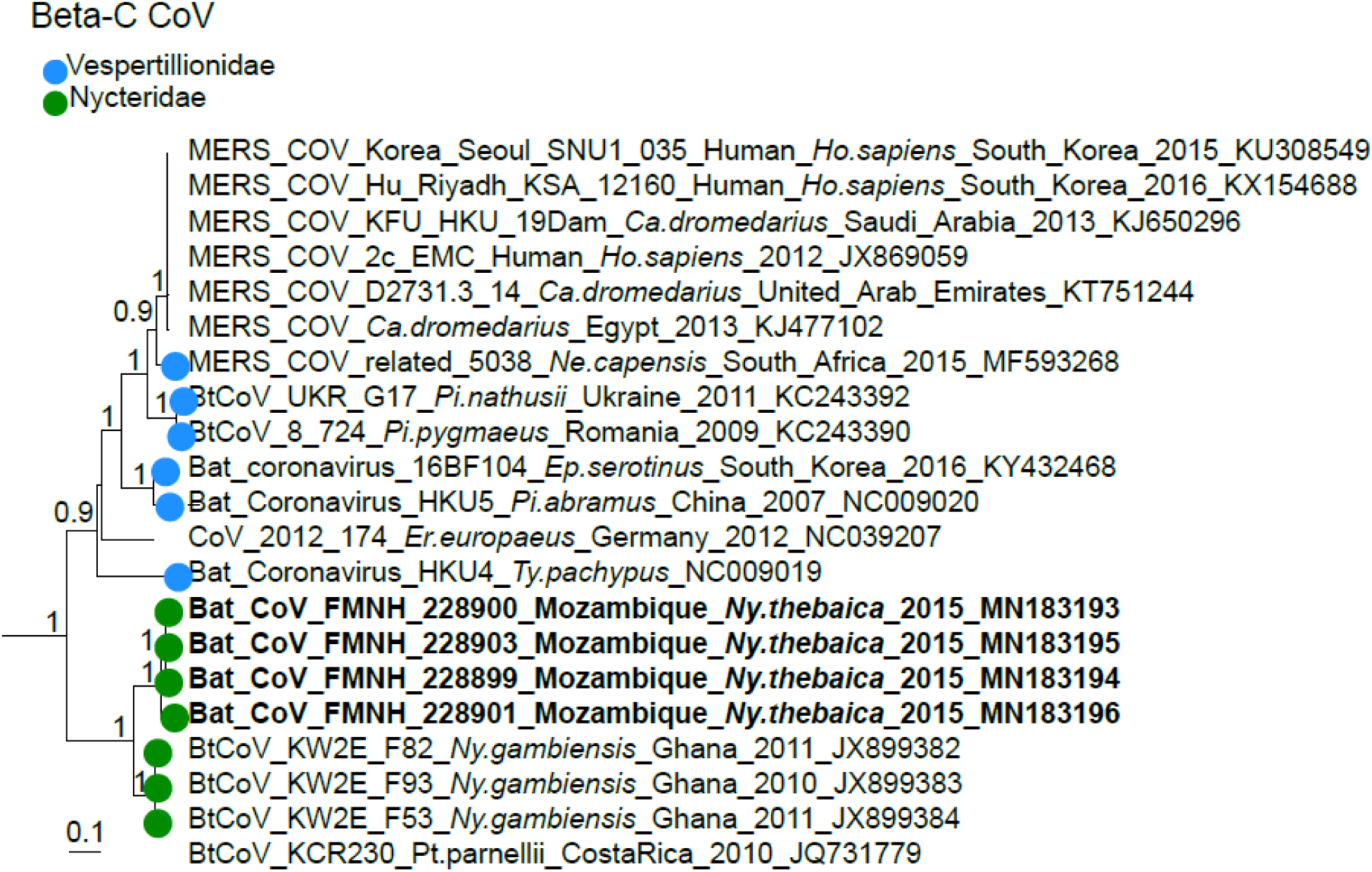
Detail of the β-C CoV clade. CoVs generated in the study are indicated in bold. This sub-tree is a zoom on β-C CoV clade from the tree depicted in Figure 3. Bootstrap values >0.7 are indicated on the tree. Scale bar indicates mean number of nucleotide substitutions per site.

**Figure 8.**
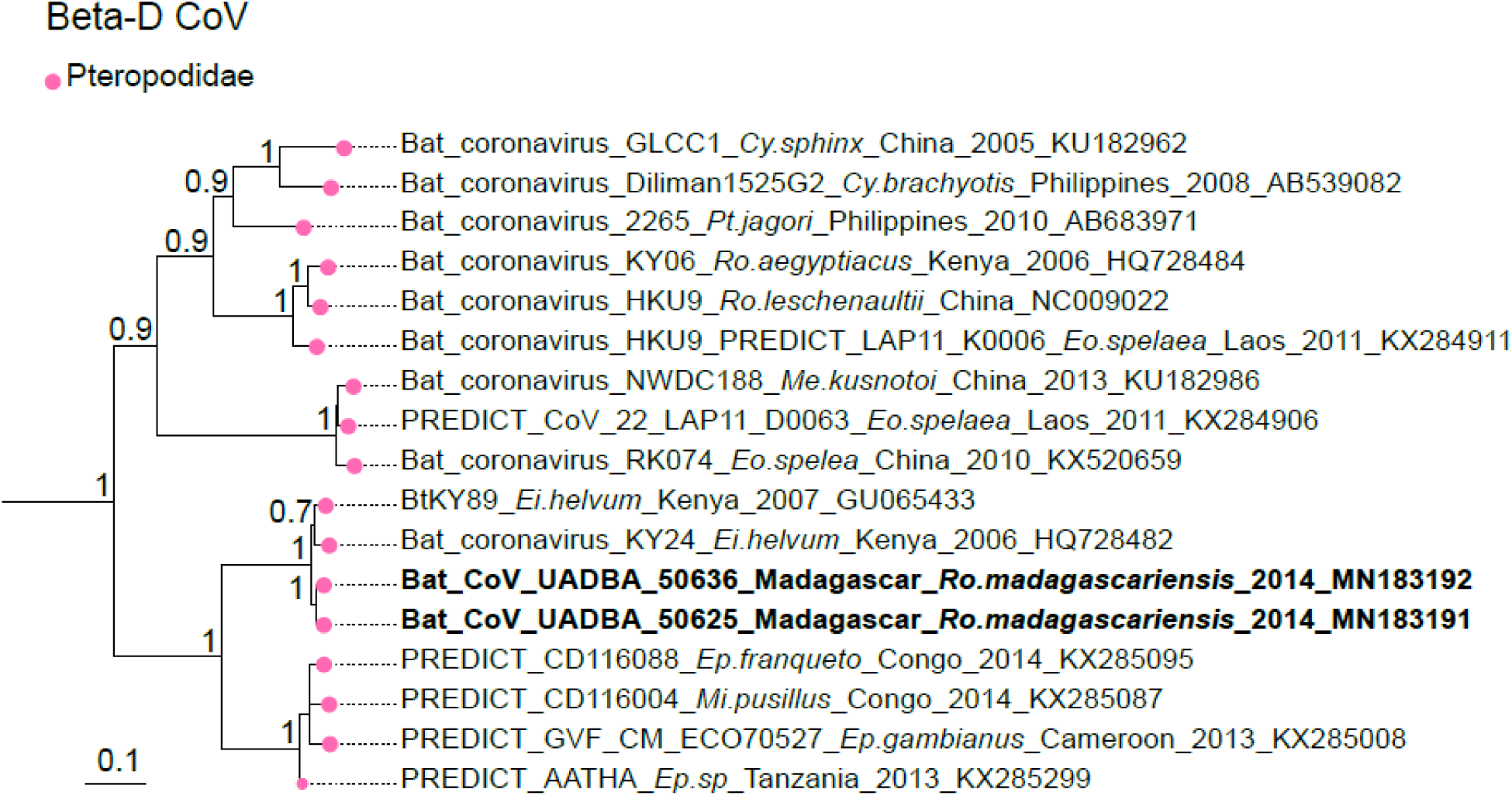
Detail of the β-D CoV. CoVs generated in the study are indicated in bold. This sub-tree is a zoom on β-D CoV clade from the tree depicted in Figure 3. Bootstrap values >0.7 are indicated on the tree. Scale bar indicates mean number of nucleotide substitutions per site.

### Co-phylogeny between bats and CoVs

Co-phylogeny tests were conducted using 11 Cyt *b* sequences obtained from the 11 CoVs positive bat species and 27 partial CoV RdRp sequences (440 bp). Results supported co-evolution between the Western Indian Ocean bats and their CoVs (ParaFitGlobal = 0.04; p = 0.001) and a high level of phylogenetic congruence (Figure 9).

**Figure 9.**
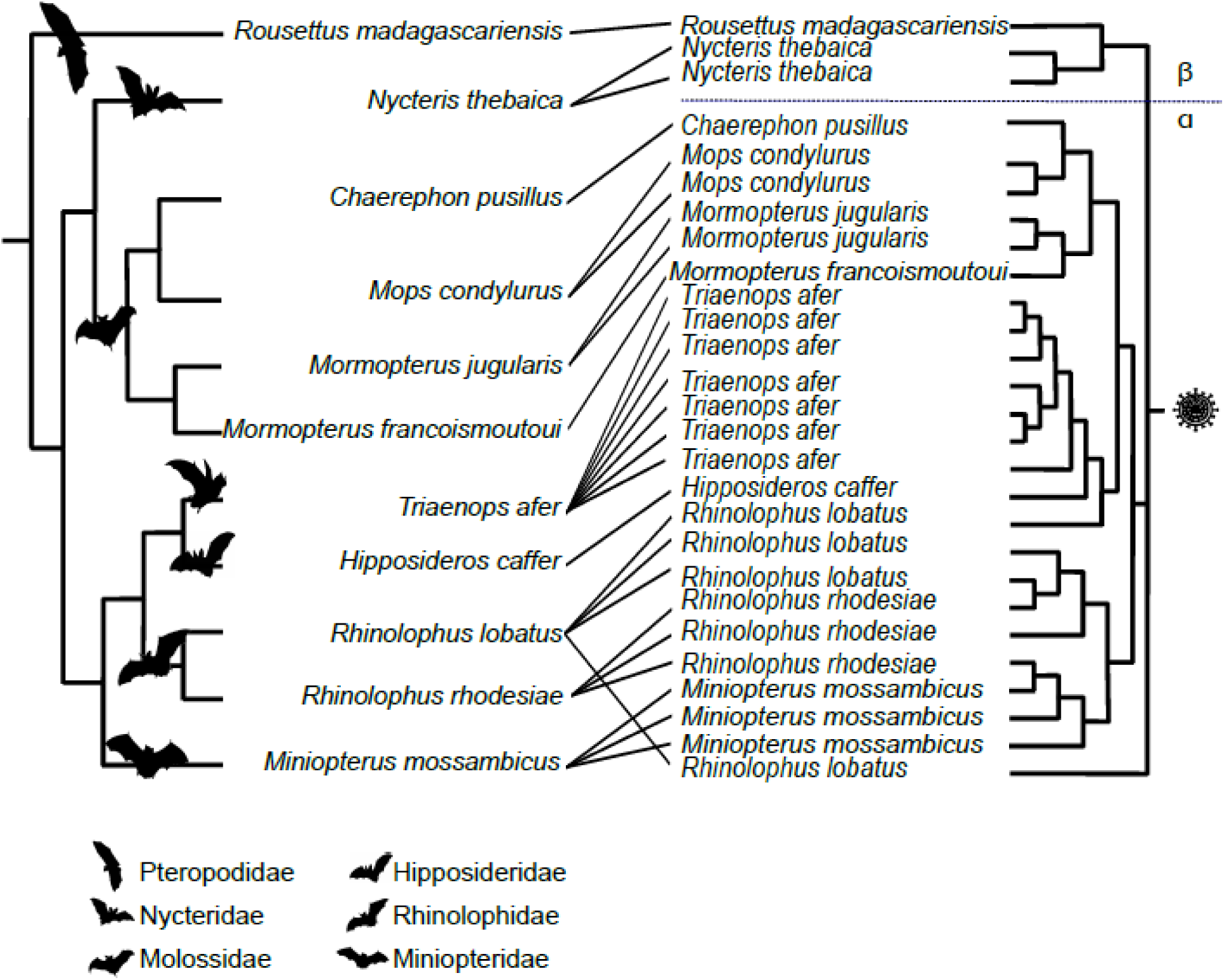
Tanglegram representing host-virus co-evolution between bats of the Western Indian Ocean and their associated CoVs. Phylogeny of bats (left) was constructed with an alignment of 11 Cyt *b* sequences of 1,030 bp by Neighbor-Joining with 1,000 bootstrap iterations. Pruned phylogeny of Western Indian Ocean bats CoVs (right) was constructed with an alignment of 27 unique sequences of 393 bp from Western Indian Ocean bats CoVs, by Neighbor-Joining with 1,000 bootstrap iterations.

## Discussion

We provide evidence for a high diversity of CoVs in bats on Western Indian Ocean islands. The overall prevalence of CoV positive bats was consistent with studies from continental Africa ^25^ and from islands in the Australasian region ^31^, although we detected significant variation in the prevalence of infected bats, according to their family, species, sampling location and season. Our study is nevertheless affected by the strong heterogeneity of bat communities in the island of the Western Indian Ocean, in particular in term of species richness. The high CoV genetic diversity detected in bats from Mozambique and Madagascar is likely to be associated with the higher bat species diversity in the African mainland and in Madagascar, has compared to small oceanic islands ^20^. In addition, CoV prevalence in bat populations may significantly vary across seasons, as found in Mozambique with higher prevalence during the wet season than in the dry season. Several studies on bat CoV have indeed shown significant variations in the temporal infection dynamic of CoV in bats, potentially associated with bat parturition ^32–34^.

Host specificity is well known for some bat CoVs subgenera ^35–37^. For example, β-C CoVs are largely associated with Vespertilionidae, whereas β-D CoVs are found mostly in Pteropodidae ^36,38^. In our study, we showed that Western Indian Ocean bats harbor phylogenetically structured CoVs, of both α-CoV and β-CoV subclades, clustering mostly by bat family. In the new CoV taxonomy based on full genomes proposed by the International Committee of Taxonomy of Viruses (ICTV), α-CoVs and β-CoVs are split in subgenera mostly based on host families ^39^, reflected in the subgenera names (e.g. Rhinacovirus for a Rhinolophidae α-CoV cluster, Minuacovirus for a Miniopteridae α-CoV cluster, Hibecovirus for an Hipposideridae β-CoV cluster). Although our classification was based on a partial sequence of the RdRp region, we identified sequences from samples belonging to four of these subgenera (Minuacovirus, Duvinacovirus, Rhinacovirus, and Nobecovirus) and three that could not be classified according to this taxonomic scheme hence representing unclassified subgenera (we propose “Molacovirus”, “Nycbecovirus”, and “Rhinacovirus2”).

A strong geographical influence on CoVs diversity, with independent evolution of CoVs on each island, was expected in our study, because of spatial isolation and endemism of the tested bat species. Anthony et al. ^38^ found that the dominant evolutionary mechanism for African CoVs was host switching. Congruence between host and viral phylogenies however suggests a strong signal for co-evolution between Western Indian Ocean bats and their associated CoVs. Geographical influence seems to occur within bat families, as for Molossidae. Endemism resulting from geographic isolation may thus have played a role in viral diversification within bat families.

Although co-evolution could be the dominant mechanism in the Western Indian Ocean, host-switching may take place in certain situations. For example, in Mozambique, we found a potential Miniopteridae α-CoV in a Rhinolophidae bat co-roosting with Miniopteridae in the same cave. These host-switching events could be favored when several bat species roost in syntopy ^40^. A similar scenario was described in Australia where Miniopteridae α-CoV was detected in Rhinolophidae bats ^31^. These infrequent host-switching events show that spillovers can happen but suggest that viral transmission is not maintained in the receiver host species. The host-virus co-evolution might thus have resulted in strong adaptation of CoVs to each bat host species. In addition, viral factors (mutation rate, recombination propensity, replication ability in the cytoplasm, changes in the ability to bind host cells), environmental factors (climate variation, habitat degradation, decrease of bat preys), and phylogenetic relatedness of host species are also critical for the viral establishment in a novel host ^41–44^. Nevertheless, apparent evidence of host switching as a dominant mechanism of CoV evolution could be an artifact of a lack of data for some potential bat hosts, leading to incomplete phylogenetic reconstructions ^38^.

Several bat CoVs we identified in Rhinonycteridae and Hipposideridae from Mozambique had between 85% and 93% nucleotide sequence similarity with NL63 Human CoVs and 229E Human CoVs, respectively. These two human viruses are widely distributed in the world and associated with mild to moderate respiratory infection in humans ^45^. Tao et al. established that the NL63 Human CoVs and 229E Human CoVs have a zoonotic recombinant origin from their most recent common ancestor, estimated to be about 1,000 years ago ^46^. During the past decade, they were both detected in bats in Kenya, and in Ghana, Gabon, Kenya, and Zimbabwe, respectively ^24,28,47,48^. Intermediate hosts are important in the spillover of CoVs, despite major knowledge gaps on the transmission routes of bat infectious agents to secondary hosts ^49^. This hypothesis has been formulated for the 229E Human CoV, with an evolutionary origin in Hipposideridae bats and with camelids as intermediate hosts ^48^. The ancient spillover of NL63 from Rhinonycteridae bats to humans might have occurred through a currently unidentified intermediate host ^28,50,51^. Because receptor recognition by viruses is the first essential cellular step to infect host cells, CoVs may have spilt over into humans from bats through an intermediate host possibly due to mutations on spike genes ^13,28^. Further investigations of CoVs in Kenyan and Mozambican livestock and hunted animals could potentially provide information on the complete evolutionary and emergence history of these two viruses before their establishment in humans.

MERS-like CoV, with high sequence similarity (>85%) to human and camel strains of MERS-CoV, have been detected in *Neoromicia capensis* in South Africa and *Pipistrellus* cf. *hesperidus* in Uganda, suggesting a possible origin of camel MERS-CoV in vespertilionid bats ^25,38,52^. This family has been widely studied, with 30% of all reported bat CoVs sequences from the past 20 years coming from vespertilionids ^53^. No members of this family were positive for CoV in our study, which may be associated with the low number of individuals tested; additional material is needed to explore potential MERS-like CoV in the Western Indian Ocean, in particular on Madagascar.

Knowledge on bat CoV ecology and epidemiology has significantly increased during the past decade. Anthony et al. estimated that there might be at least 3,204 bat CoVs worldwide ^38^; however, direct bat-to-human transmission has not been demonstrated so far. As for most emerging zoonoses, CoV spillover and emergence may be associated to human activities and ecosystem changes such as habitat fragmentation, agricultural intensification and bushmeat consumption. The role of bats as epidemiological reservoir of infectious agents needs to be balanced with such human driven modifications on ecosystem functioning, in order to properly assess bat-borne CoV emergence risks.

## Materials and methods

### Origin of the tested samples

Samples obtained from vouchered bat specimens during previous studies in Mozambique (February and May 2015), Mayotte (November to December 2014), Reunion Island (February 2015), Seychelles (February to March 2014), Mauritius (November 2012) and Madagascar (October to November 2014) were tested ^54–57^ (Supplementary Information). We also collected additional swab samples from several synanthropic bat species on Madagascar, in January 2018 (Supplementary Information). Details on sample types, bat families, species, and locations are provided in Supplementary Table S3.

### Ethical statement

The ethical terms of these research protocols were approved by the CYROI Institutional Animal Care and Use Committee (Comité d’Ethique du CYROI no.114, IACUC certified by the French Ministry of Higher Education, of Research and Innovation). All protocols strictly followed the terms of research permits and regulations for the handling of wild mammals and were approved by licencing authorities (Supplementary Information).

### Molecular detection

RNA was extracted from 140 μL of each sample using the QIAamp Viral RNA mini kit (QIAGEN, Valencia, California, USA), and eluted in 60 μL of Qiagen AVE elution buffer. For bat organs, approximately 1 mm^3^ of tissue (either lungs or intestines) was placed in 750 µL of DMEM medium and homogenized in a TissueLyser II (Qiagen, Hilden, Germany) for 2 min at 25 Hz using 3 mm tungsten beads, prior to the RNA extraction. Reverse transcription was performed on 10 μL of RNA using the ProtoScript II Reverse Transcriptase and Random Primer 6 (New England BioLabs, Ipswich, MA, USA) under the following thermal conditions: 70 °C for 5 min, 25 °C for 10 min, 42 °C for 50 min, and 65 °C for 20 min ^58^. cDNAs were tested for the presence of the RNA-dependent RNA-polymerase (RdRp) gene using a multi-probe Real-Time PCR ^59^. The primer set with Locked Nucleic Acids (LNA; underlined position in probe sequences) was purchased from Eurogentec (Seraing, Belgium): 11-FW: 5’-TGA-TGA-TGS-NGT-TGT-NTG-YTA-YAA-3’ and 13-RV: 5’-GCA-TWG-TRT-GYT-GNG-ARC-ARA-ATT-C-3’. Three probes were used: probe I (ROX): 5’-TTG-<u>T</u>AT-<u>T</u>AT-<u>C</u>AG-A<u>A</u>T-G<u>G</u>Y-G<u>T</u>S-<u>T</u>TY-A<u>T</u>-3’, probe II (FAM): 5’-TGT-GTT-CAT-GTC-WGA-RGC-WAA-ATG-TT-3’, and probe III (HEX): 5’-T<u>C</u>T-A<u>A</u>R-<u>T</u>GT-<u>T</u>G<u>G-</u>GTD-<u>G</u>A-3’. Real-Time PCR was performed with ABsolute Blue QPCR Mix low ROX 1X (Thermo Fisher Scientific, Waltham, MA, USA) and 2.5 µL of cDNA under the following thermal conditions: 95 °C for 15 min, 95 °C for 30 s, touchdowns from 56 °C to 50°C for 1 min and 50 cycles with 95 °C for 30 s and 50 °C for 1 min in a CFX96 Touch Real-Time PCR Detection System (Bio-Rad, Hercules, CA, USA).

Because of the limited size of sequences generated from the Real-Time PCR, a second PCR targeting 440 bp of the RdRp gene was performed with 5 µL of cDNA of each positive sample, with the following primer set: IN-6: 5’-GGT-TGG-GAC-TAT-CCT-AAG-TGT-GA-3’ and IN-7: 5’-CCA-TCA-TCA-GAT-AGA-ATC-ATC-ATA-3’ ^60^. PCRs were performed with the GoTaq G2 Hot Start Green Master Mix (Promega, Madison, WI, USA) in an Applied Biosystems 2720 Thermal Cycler (Thermo Fisher Scientific, Waltham, MA, USA), under the following thermal conditions: 95 °C for 2 min, 45 cycles with 95 °C for 1 min, 54 °C for 1 min, 72°C for 1 min, and a final elongation step at 72°C for 10 min. After electrophoresis in a 1.5% agarose gel stained with 2% GelRed (Biotium, Hayward, CA, USA), amplicons of the expected size were sequenced on both strands by Genoscreen (Lille, France). All generated sequences were deposited in GenBank under the accession numbers MN183146 to MN183273.

### Statistical analysis

We have performed Pearson χ^2^ tests on all samples (1,013 bats) to explore the effect of (i) location, (ii) bat family, and (iii) bat species on the detection of coronavirus RNA. Two sampling campaigns, at two different season, in the same location, were available for Mozambique. We thus investigated the effect of the sampling season, between the wet (February) and dry (May) season, on CoV detection in Mozambique in 2015 (264 bats). Analyses were conducted with R v3.5.1 software ^61^.

### Phylogenetic analyses

Sequences obtained with the second PCR system ^60^ were edited with the Chromas Lite Software package version 2.6.4 ^62^. We explored CoV diversity of the sequences with pairwise identity values obtained from *seqidentity* function in R *bio3d* package v2.3-4 ^63^ and identified the most similar CoV RdRp sequences referenced in GenBank using BLASTN 2.2.29+. An alignment was then generated using the 51 nucleotide sequences obtained in this study and 151 reference CoV sequences representing a large diversity of hosts and geographic origins (Europe, Asia, Oceania, America and Africa), with CLC Sequence viewer 8.0 Software (CLC Bio, Aarhus, Denmark). A phylogenetic tree was obtained by maximum likelihood using MEGA Software v10.0.4 ^64^, with 1,000 bootstrap iterations, and with the best evolutionary model for our dataset as selected by modelgenerator v0.85 ^65^.

Host-virus associations were investigated using the phylogeny of Western Indian Ocean bats and their associated CoVs. Bat phylogeny was generated from an alignment of 1,030 bp of mitochondrial Cytochrome *b* (Cyt *b*) gene sequences (Supplementary Table S4), for each CoV positive bat species. Finally, bat and pruned CoV phylogenies based on each 393 bp RdRp unique sequence fragment were generated by Neighbor-Joining with 1,000 bootstrap iterations, using CLC Sequence viewer 8.0 Software (CLC Bio, Aarhus, Denmark)^66^. Phylogenetic congruence was tested to assess the significance of the coevolutionary signal between bat host species and CoVs sequences, using *ParaFit* with 999 permutations in the *ape* package v5.0 in R 3.5.1 ^67,68^. Tanglegram representations of the co-phylogeny were visualized using the Jane software v4.01 ^69^.

## Acknowledgments

We are very grateful to S. Muradrasoli for recommendations on the optimization of the multi-probe PCR protocol as well as for providing PCR controls and to C. Cordonin for providing bat Cyt *b* sequences. L. Biscornet, C. Dionisio, L. Domergue, M. Dietrich, T. Mbohoahy, T. Nekena, J. Rakotoarivelo, M. Rakotomanga, C. F. Rakotondramanana, and A. Randrenjarison are thanked for their assistance in the field. We also thank S. Bos and A. Hoarau for their help in the laboratory, and K. Dellagi and H. Pascalis for the development and the management of the ‘partenariat Mozambique-Réunion dans la recherche en santé: pour une approche intégrée d’étude des maladies infectieuses à risque épidémique (MoZaR)’ research program.

## Data availability

DNA sequences: Genbank accessions MN183146 to MN183273

