## Supplementary Information for "Bat coronavirus phylogeography in the Western Indian Ocean"

***Research permits for the collection of biological materials***

### Reunion Island

Samples were collected as part of a previous study on lyssavirus infection in bats in the Indian Ocean (1), under a research permit delivered by the Préfecture de La Réunion: “Arrêté préfectoral du 11 Février 2013 and Arrêté préfectoral du 11 Septembre 2014 (N°2014-07)”.

### Mauritius

Samples were collected as part of a previous study on lyssavirus infection in bats in the Indian Ocean (1), under a memorandum of agreement for the supply of biological material by Government of Mauritius (delivered by the National Park and Conservation Service for authorization of Mauritius), signed 17 December 2010 and 09 January 2013. CITES permit from the Mauritian national authority was issued for tissue export (permit MU120933) to Reunion Island.

### Mayotte

Samples were collected as part of a previous study on lyssavirus infection in bats in the Indian Ocean (1), under a research permit delivered by the “Préfecture de Mayotte: Arrêté N°158/DEAL/SEPR/2014”.

### Madagascar

Samples were collected as part of previous studies on infectious agents in Malagasy wildlife (1–4), under the following research permits delivered by the “Direction du Système des Aires Protégées and Direction Générale de l’Environnement et des Forêts”; Madagascar National Parks: 350/10/MEF/SG/DGF/DCB.SAP/SCB, 032/12/MEF/SG/DGF/ DCB.SAP/SCBSE, 067/12/MEF/SG/DGF/DCB.SAP/SCBSE, 194/12/ MEF/SG/DGF/DCB.SAP/SCB, N◦283/11/MEF/SG/DGF/DCB.SAP/SCB, N◦077/12/MEF/SG/DGF/DCB.SAP/SCBSE, 238/14/MEEF/SG/ DGF/DCB.SAP/SCB and 268/14/MEEF/SG/DGF/DCB.SAP/SCB. A CITES permit from the Malagasy national authority was issued for tissue export (permit 243C-EA06/MG12) to Reunion island.

### Seychelles (Mahe)

Samples were collected as part of a previous study on lyssavirus infection in bats in the Indian Ocean (1), under the following research permit delivered by the Direction of Wildlife, Trade and Conservation Section and the Ministry of Environment and Energy of Republic of Seychelles: Export permits Agreement of 5 March 2014. CITES permit from the Republic of Seychelles was issued for export (permit N°1772) to Reunion Island.

### Mozambique (Inhassoro district, Inhambane province)

Samples were collected as part of a previous study on astrovirus infection in bats in the Indian Ocean (5), under a research permit delivered by the Ministry of Health (N°S/N/SDI/0233/15).

#### *Type of collected samples*

Organs were collected before 2014, stored in liquid nitrogen in the field and then at −80°C upon arrival at the laboratory (Appendix Table 1). From 2014 onward, feces, rectal, and oral swabs were collected (Appendix Table 1) and placed in 1mL brain heart infusion medium (Conda, Madrid, Spain) supplemented with penicillin G (1,000 units/mL), streptomycin (1 mg/mL), kanamycin (0.5 mg/mL), gentamicin (0.25 mg/mL) and amphotericin B (0.025 mg/mL). These samples were also stored in liquid nitrogen in the field and at −80°C upon arrival at the laboratory (Appendix Table 1).

### Supplementary tables

**Table S1.** Number of positive samples and sequences generated with the Real-Time PCR detection system, and the classic PCR system used to generate sequences for phylogenetic analyses.

| | | | | Detection system | | | Phylogeny system | |
|---|---|---|---|---|---|---|---|---|
| **Location** | **Sampling year** | **Bat family** | **Bat species** | **N tested** | **N detected** | **N sequences** | **N tested** | **N sequences** |
| **Madagascar** | **2012** | **Molossidae** | *Mops midas* | 21 | 4 | 4 | 4 | 0 |
| | | **Rhinonycteridae** | *Triaenops menamena* | 21 | 2 | 0 | 2 | 0 |
| | **2013** | **Molossidae** | *Mormopterus jugularis* | 63 | 10 | 7 | 10 | 5 |
| | | **Pteropodidae** | *Rousettus madagascariensis* | 45 | 6 | 6 | 6 | 2 |
| | **2014** | **Rhinonycteridae** | *Triaenops menamena* | 13 | 2 | 2 | 2 | 0 |
| **Mayotte** | **2014** | **Molossidae** | *Chaerephon sp* | 4 | 1 | 1 | 1 | 1 |
| | | | *Chaerephon pusillus* | 61 | 7 | 7 | 7 | 5 |
| **Mozambique** | **2015** | **Hipposideridae** | *Hipposideros caffer* | 59 | 10 | 6 | 10 | 3 |
| | | **Miniopteridae** | *Miniopterus mossambicus* | 21 | 4 | 4 | 4 | 4 |
| | | **Molossidae** | *Mops condylurus* | 54 | 11 | 11 | 11 | 4 |
| | | **Nycteridae** | *Nycteris thebaica* | 14 | 4 | 3 | 4 | 4 |
| | | **Rhinolophidae** | *Rhinolophus lobatus* | 9 | 6 | 6 | 6 | 4 |
| | | | *Rhinolophus rhodesiae* | 30 | 9 | 9 | 9 | 9 |
| | | | *Rhinolophus sp* | 2 | 1 | 1 | 1 | 1 |
| | | **Rhinonycteridae** | *Triaenops afer* | 51 | 9 | 8 | 9 | 8 |
| **Reunion** | **2015** | **Molossidae** | *Mormopterus francoismoutoui* | 50 | 2 | 2 | 2 | 1 |
| | | | | | 88 | 77 | 88 | 51 |

**Table S2.** Nucleotide sequence similarity obtained with BLAST.

| ID | species | % identity | GENBANK Sequence | Accession GENBANK ID |
|---|---|---|---|---|
| MAY004 | *Chaerephon pusillus* | 92% | Chaerephon bat coronavirus/Kenya/KY22/2006 polyprotein (ORF1ab) gene | HQ728486 |
| MAY015 | *Chaerephon pusillus* | 92% | Chaerephon bat coronavirus/Kenya/KY22/2006 polyprotein (ORF1ab) gene | HQ728486 |
| MAY025 | *Chaerephon pusillus* | 92% | Chaerephon bat coronavirus/Kenya/KY22/2006 polyprotein (ORF1ab) gene | HQ728486 |
| MAY027 | *Chaerephon pusillus* | 92% | Chaerephon bat coronavirus/Kenya/KY22/2006 polyprotein (ORF1ab) gene | HQ728486 |
| MAY033 | *Chaerephon pusillus* | 92% | Chaerephon bat coronavirus/Kenya/KY22/2006 polyprotein (ORF1ab) gene | HQ728486 |
| MAY051 | *Chaerephon sp* | 92% | Chaerephon bat coronavirus/Kenya/KY22/2006 polyprotein (ORF1ab) gene | HQ728486 |
| RB369 | *Mormopterus francoismoutoui* | 90% | Chaerephon bat coronavirus/Kenya/KY22/2006 polyprotein (ORF1ab) gene | HQ728486 |
| SMG18218 | *Mormopterus jugularis* | 90% | Chaerephon bat coronavirus/Kenya/KY22/2006 polyprotein (ORF1ab) gene | HQ728486 |
| SMG18223 | *Mormopterus jugularis* | 90% | Chaerephon bat coronavirus/Kenya/KY22/2006 polyprotein (ORF1ab) gene | HQ728486 |
| SMG18225 | *Mormopterus jugularis* | 90% | Chaerephon bat coronavirus/Kenya/KY22/2006 polyprotein (ORF1ab) gene | HQ728486 |
| SMG18231 | *Mormopterus jugularis* | 90% | Chaerephon bat coronavirus/Kenya/KY22/2006 polyprotein (ORF1ab) gene | HQ728486 |
| SMG18238 | *Mormopterus jugularis* | 90% | Chaerephon bat coronavirus/Kenya/KY22/2006 polyprotein (ORF1ab) gene | HQ728486 |
| SMG18866 | *Rousettus madagascariensis* | 98% | Kenya bat coronavirus BtKY89 RNA-dependent RNA polymerase gene, partial cds | GU065433 |

| | | | | |
|---|---|---|---|---|
| SMG18930 | *Rousettus madagascariensis* | 98% | Kenya bat coronavirus BtKY89 RNA-dependent RNA polymerase gene, partial cds | GU065433 |
| SMG18951 | *Hipposideros caffer* | 96% | 229E-related bat coronavirus strain BtKY229E-8 | KY073748 |
| SMG18958 | *Triaenops afer* | 98% | NL63-related bat coronavirus strain BtKYNL63-15, complete genome | KY073746 |
| SMG18970 | *Nycteris thebaica* | 88% | Betacoronavirus BtCoV/KW2E-F93/Nyc_spec/GHA/2010 RNA-dependent RNA polymerase gene, partial cds; and spike glycoprotein gene, complete cds | JX899383 |
| SMG18971 | *Nycteris thebaica* | 88% | Betacoronavirus BtCoV/KW2E-F93/Nyc_spec/GHA/2010 RNA-dependent RNA polymerase gene, partial cds; and spike glycoprotein gene, complete cds | JX899383 |
| SMG18972 | *Nycteris thebaica* | 88% | Betacoronavirus BtCoV/KW2E-F93/Nyc_spec/GHA/2010 RNA-dependent RNA polymerase gene, partial cds; and spike glycoprotein gene, complete cds | JX899383 |
| SMG18987 | *Hipposideros caffer* | 96% | 229E-related bat coronavirus strain BtKY229E-8, complete genome | KY073748 |
| SMG18990 | *Rhinolophus lobatus* | 87% | Kenya bat coronavirus BtKY83 RNA-dependent RNA polymerase gene, partial cds | GU065427 |
| SMG18991 | *Rhinolophus lobatus* | 87% | Kenya bat coronavirus BtKY83 RNA-dependent RNA polymerase gene, partial cds | GU065427 |
| SMG18992 | *Rhinolophus lobatus* | 87% | Kenya bat coronavirus BtKY83 RNA-dependent RNA polymerase gene, partial cds | GU065427 |
| SMG18993 | *Triaenops afer* | 98% | NL63-related bat coronavirus strain BtKYNL63-15, complete genome | KY073746 |
| SMG18996 | *Triaenops afer* | 98% | NL63-related bat coronavirus strain BtKYNL63-15, complete genome | KY073746 |
| SMG18997 | *Triaenops afer* | 98% | NL63-related bat coronavirus strain BtKYNL63-15, complete genome | KY073746 |
| SMG19002 | *Hipposideros caffer* | 96% | 229E-related bat coronavirus strain BtKY229E-8, complete genome | KY073748 |

| | | | | |
|---|---|---|---|---|
| SMG19004 | *Miniopterus mossambicus* | 98% | Miniopterus bat coronavirus/Kenya/KY27/2006 polyprotein (ORF1ab) gene | HQ728484 |
| SMG19006 | *Miniopterus mossambicus* | 97% | Miniopterus bat coronavirus/Kenya/KY27/2006 polyprotein (ORF1ab) gene | HQ728484 |
| SMG19013 | *Miniopterus mossambicus* | 98% | Miniopterus bat coronavirus/Kenya/KY27/2006 polyprotein (ORF1ab) gene | HQ728484 |
| SMG19015 | *Miniopterus mossambicus* | 98% | Miniopterus bat coronavirus/Kenya/KY27/2006 polyprotein (ORF1ab) gene | HQ728484 |
| SMG19020 | *Rhinolophus rhodesiae* | 87% | Kenya bat coronavirus BtKY83 RNA-dependent RNA polymerase gene, partial cds | GU065427 |
| SMG19024 | *Rhinolophus rhodesiae* | 98% | Miniopterus bat coronavirus/Kenya/KY27/2006 polyprotein (ORF1ab) gene | HQ728484 |
| SMG19026 | *Rhinolophus sp* | 87% | Kenya bat coronavirus BtKY83 RNA-dependent RNA polymerase gene, partial cds | GU065427 |
| SMG19027 | *Rhinolophus rhodesiae* | 87% | Kenya bat coronavirus BtKY83 RNA-dependent RNA polymerase gene, partial cds | GU065427 |
| SMG19028 | *Rhinolophus rhodesiae* | 87% | Kenya bat coronavirus BtKY83 RNA-dependent RNA polymerase gene, partial cds | GU065427 |
| SMG19029 | *Rhinolophus lobatus* | 82% | Rhinolophus bat coronavirus BtCoV/4307-2 RNA-dependent RNA polymerase (RdRp) gene, partial cds | KP876528 |
| SMG19030 | *Rhinolophus rhodesiae* | 87% | Kenya bat coronavirus BtKY83 RNA-dependent RNA polymerase gene, partial cds | GU065427 |
| SMG19031 | *Rhinolophus rhodesiae* | 87% | Kenya bat coronavirus BtKY83 RNA-dependent RNA polymerase gene, partial cds | GU065427 |
| SMG19032 | *Rhinolophus rhodesiae* | 87% | Kenya bat coronavirus BtKY83 RNA-dependent RNA polymerase gene, partial cds | GU065427 |
| SMG19037 | *Rhinolophus rhodesiae* | 87% | Kenya bat coronavirus BtKY83 RNA-dependent RNA polymerase gene, partial cds | GU065427 |
| SMG19038 | *Rhinolophus rhodesiae* | 82% | Rhinolophus bat coronavirus BtCoV/4307-2 RNA-dependent RNA polymerase (RdRp) gene, partial cds | KP876528 |
| SMG19141 | *Mops condylurus* | 98% | Chaerephon bat coronavirus/Kenya/KY22/2006 polyprotein (ORF1ab) gene | HQ728486 |

| | | | | |
|---|---|---|---|---|
| SMG19154 | *Triaenops afer* | 98% | NL63-related bat coronavirus strain BtKYNL63-15, complete genome | KY073746 |
| SMG19174 | *Triaenops afer* | 98% | NL63-related bat coronavirus strain BtKYNL63-15, complete genome | KY073746 |
| SMG19183 | *Triaenops afer* | 98% | NL63-related bat coronavirus strain BtKYNL63-15, complete genome | KY073746 |
| SMG19185 | *Triaenops afer* | 98% | NL63-related bat coronavirus strain BtKYNL63-9a | KY073744 |
| SMG19207 | *Mops condylurus* | 98% | Chaerephon bat coronavirus/Kenya/KY22/2006 polyprotein (ORF1ab) gene | HQ728486 |
| SMG19210 | *Mops condylurus* | 98% | Chaerephon bat coronavirus/Kenya/KY22/2006 polyprotein (ORF1ab) gene | HQ728486 |
| SMG19217 | *Mops condylurus* | 98% | Chaerephon bat coronavirus/Kenya/KY22/2006 polyprotein (ORF1ab) gene | HQ728486 |

**Table S3.** Number of tested and positive samples, per location, bat family, species, samples types, and collection year.

| Location | Bat family | Bat species | Collection Year | N tested | N tested per type of samples | | | | N positive (%) | | | |
|---|---|---|---|---|---|---|---|---|---|---|---|---|
| | | | | | Organs | | Feces | Swabs | Organs | | Feces | Swabs |
| | | | | | Lung | Intestine | | | Lung | Intestine | | |
| Madagascar | Hipposideridae | *Macronycteris commersoni* | 2012 | 19 | | 19 | | | | | | |
| | Miniopteridae | *Miniopterus manavi* | 2012 | 19 | | 19 | | | | | | |
| | | *Miniopterus gleni* | 2012 | 14 | | 14 | | | | | | |
| | | | 2014 | 2 | | | | 2 | | | | |
| | | *Miniopterus griveaudi* | 2014 | 28 | | | | 28 | | | | |
| | | *Miniopterus mahafaliensis* | 2012 | 8 | | 8 | | | | | | |
| | | *Miniopterus sororculus* | 2012 | 8 | | 8 | | | | | | |
| | Molossidae | *Chaerephon atsinanana* | 2013 | 29 | | 29 | | | | | | |
| | | *Chaerephon leucogaster* | 2018 | 45 | | | | 45 | | | | |
| | | *Mops leucostigma* | 2018 | 94 | | | | 94 | | | | |
| | | ***Mops midas*** | 2012 | 19 | | 19 | | | | 4 (21.1) | | |
| | | | 2018 | 2 | | | | 2 | | | | |
| | | ***Mormopterus jugularis*** | 2013 | 22 | | 22 | | | | 10 (45.5) | | |
| | | | 2014 | 41 | | | | 41 | | | | |
| | | *Otomops madagascariensis* | 2012 | 12 | | 12 | | | | | | |
| | | | 2014 | 6 | | | | 6 | | | | |
| | Pteropodidae | ***Rousettus madagascariensis*** | 2014 | 45 | | | | 45 | | | | 6 (13.3) |
| | Rhinonycteridae | *Paratriaenops furculus* | 2014 | 32 | | | | 32 | | | | |
| | | ***Triaenops menamena*** | 2012 | 21 | | 21 | | | | 2 (9.5) | | |
| | | | 2014 | 13 | | | | 13 | | | | 2 (15.4) |
| | Vespertilionidae | *Myotis goudoti* | 2014 | 17 | | | | 17 | | | | |
| | | *Neoromicia malagasyensis* | 2013 | 2 | | 2 | | | | | | |

| | | | | | | | | | | | | |
|---|---|---|---|---|---|---|---|---|---|---|---|---|
| | | *Neoromicia matroka* | 2013 | 4 | | 4 | | | | | | |
| | | *Pipistrellus hesperidus* | 2014 | 5 | | | | 5 | | | | |
| Mauritius | Molossidae | *Momopterus acetabulosus* | 2012 | 6 | | 6 | | | | | | |
| | Pteropodidae | *Pteropus niger* | 2012 | 48 | 48 | | | | | | | |
| | Emballonuridae | *Taphosius mauritianus* | 2014 | 2 | | | | 2 | | | | |
| Mayotte | Molossidae | ***Chaerephon sp*** | 2014 | 4 | | | 4 | | | | 1 (25.0) | |
| | | ***Chaerephon pusillus*** | 2014 | 60 | | | 60 | | | | 7 (11.6) | |
| | Pteropodidae | *Pteropus seychellensis comorensis* | 2014 | 22 | | | | 22 | | | | |
| Mozambique | Hipposideridae | ***Hipposideros caffer*** | 2015 | 59 | | | | 59 | | | | 10 (16.9) |
| | Miniopteridae | ***Miniopterus mossambicus*** | 2015 | 21 | | | | 21 | | | | 4 (19.0) |
| | Molossidae | ***Mops condylurus*** | 2015 | 54 | | | | 54 | | | | 11 (20.4) |
| | Nycteridae | ***Nycteris thebaica*** | 2015 | 14 | | | | 14 | | | | 4 (28.6) |
| | Rhinolophidae | ***Rhinolophus lobatus*** | 2015 | 9 | | | | 9 | | | | 6 (66.7) |
| | | *Rhinolophus mossambicus* | 2015 | 20 | | | | 20 | | | | |
| | | ***Rhinolophus rhodesiae*** | 2015 | 30 | | | | 30 | | | | 9 (30.0) |
| | | ***Rhinolophus sp*** | 2015 | 2 | | | | 2 | | | | 1 (50.0) |
| | Rhinonycteridae | ***Triaenops afer*** | 2015 | 51 | | | | 51 | | | | 9 (17.7) |
| | Vespertilionidae | *Neoromicia nana* | 2015 | 2 | | | | 2 | | | | |
| | | *Scotophilus viridis* | 2015 | 2 | | | | 2 | | | | |
| Reunion | Molossidae | ***Mormopterus francoismoutoui*** | 2015 | 50 | | | | 50 | | | | 2 (4.0) |

| | | | | | | | | | | | |
|---|---|---|---|---|---|---|---|---|---|---|---|
| Seychelles | Pteropodidae | *Pteropus seychellensis seychellensis* | 2014 | 50 | 50 | | | | | | |
| | | ***TOTAL*** | | **1013** | 98 | 183 | 64 | 668 | 16 | 8 | 64 |

**Table S4.** Cytochrome *b* gene sequences accession numbers from western Indian Ocean bats. *: sequences generated in this study.

| Bat family | Bat species | Cytb GenBank accession ID |
|---|---|---|
| Hipposideridae | *Hipposideros caffer* | FJ347980 |
| Miniopteridae | *Miniopterus mossambicus* | MN273501* |
| Molossidae | *Chaerephon pusillus* | GQ489154 |
| | *Mops condylurus* | GQ489181 |
| | *Mormopterus francoismoutoui* | MN273499* |
| | *Mormopterus jugularis* | KR606332 |
| Nycteridae | *Nycteris thebaica* | MN273500* |
| Pteropodidae | *Rousettus madagascariensis* | GU228727 |
| Rhinolophidae | *Rhinolophus lobatus* | MN273497* |
| | *Rhinolophus rhodesiae* | MN273498* |
| Rhinonycteridae | *Triaenops afer* | EU798748 |

**Supplementary figures**

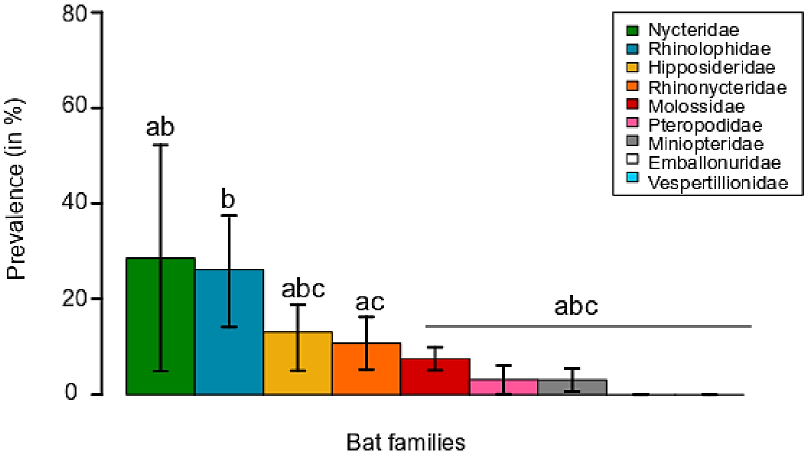


**Figure S1.** Mean CoV prevalence (± 95% confidence interval) as function of the bat family. Letters a–c above the bars refer to significantly different averages based upon a Pairwise test. Bars can have more than one letter to reflect the "overlap" between them.

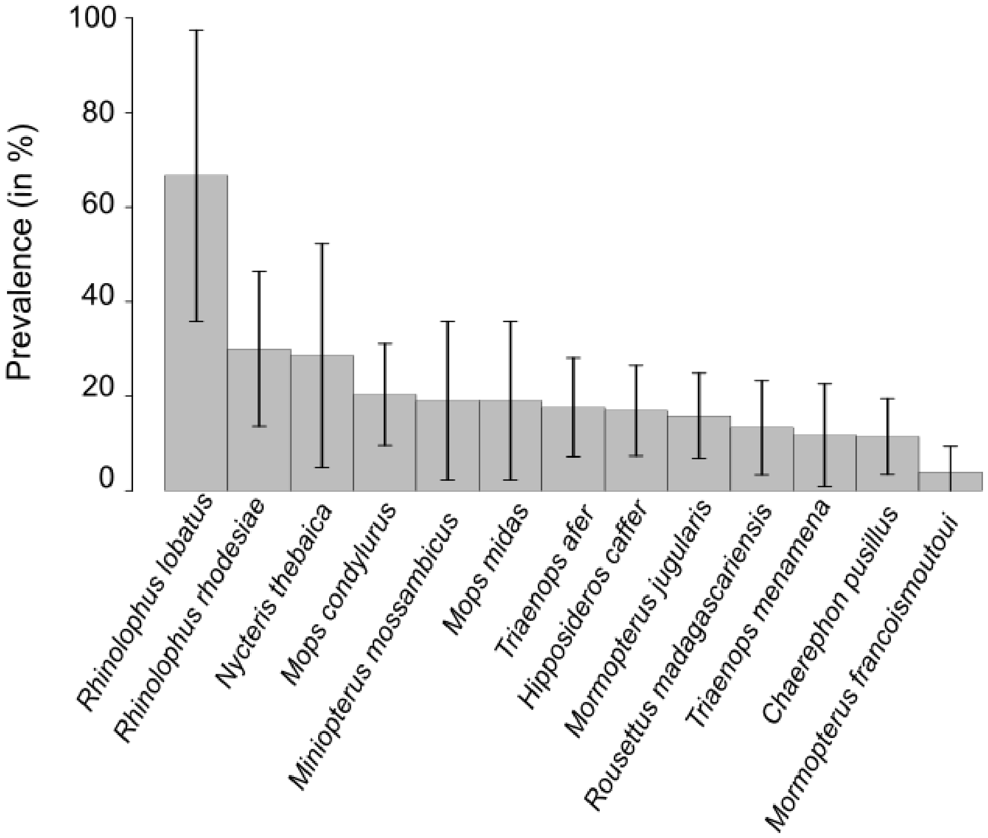


**Figure S2**. Mean CoV prevalence (mean ± 95% confidence interval) as function of the bat species.

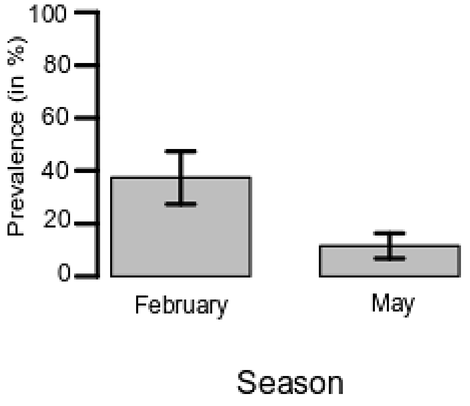


**Figure S3**. Mean CoV prevalence (mean ± 95% confidence interval) as function of the bat sampling season in Mozambique.
